# Hippocampal representations switch from errors to predictions during acquisition of predictive associations

**DOI:** 10.1101/2021.09.21.461228

**Authors:** Fraser Aitken, Peter Kok

## Abstract

We constantly exploit the statistical regularities in our environment to help guide our perception. The hippocampus has been suggested to play a pivotal role in both learning environmental statistics, as well as exploiting them to generate perceptual predictions. However, it is unclear how the hippocampus balances encoding new predictive associations with the retrieval of existing ones. Here, we present the results of two high resolution human fMRI studies (N=24 for both experiments) directly investigating this. Participants were exposed to auditory cues that predicted the identity of an upcoming visual shape (with 75% validity). Using multivoxel decoding analysis, we found that the hippocampus initially preferentially represented unexpected shapes (i.e., those that violated the cue regularities), but later switched to representing the cue-predicted shape regardless of which was actually presented. These findings demonstrate that the hippocampus in involved both acquiring and exploiting predictive associations, and switches between these modes depending on whether learning is ongoing or complete.

## Introduction

We constantly exploit the statistical regularities in our environment to help guide our perception^1^. For instance, hearing a particular jingle will prime our sensory systems for the sight (and taste!) of ice cream. But how does the brain acquire and exploit knowledge about such regularities in a changing environment?

The hippocampus has been suggested to play a pivotal role in this process. That is, the hippocampus has been shown to be involved in learning novel associations between arbitrary stimuli^2–9^, especially when stimuli are discontiguous in space and time^10–12^, as is the case for many predictive contextual cues. In fact, learning of such relationships is strongly impaired when the hippocampus is damaged^13–18^. At the same time, the hippocampus has also been suggested to play a role in exploiting such predictive associations once learning is complete^1,19,20^. Specifically, one of the main computational functions of the hippocampus is to retrieve associated items from memory based on partial information, a process known as pattern completion^21–23^. This function has mostly been studied in the context of memory recall, but is also ideally suited for retrieving perceptual predictions based on contextual cues^5,24–28^.

This raises the question of how the hippocampus balances encoding of new associations with the retrieval of existing ones^29^. One way to achieve this would be to emphasise prediction errors when an environment is novel, since these can serve to update one’s internal model of the world^30^bar. On the other hand, once an environment (and its statistical regularities) have become familiar, prediction errors may be downweighted and predictions (i.e., retrieval of existing associations) may dominate. Indeed, many previous studies have reported prediction error signals (i.e., a response evoked by mismatch between representations retrieved from memory and current sensory inputs) in the hippocampus^31–36^, while others have instead revealed prediction signals (i.e. a representation of a predicted stimulus, regardless of whether it is actually presented)^5,26,27,37^. Potentially, this seeming contradiction may arise from the fact that mismatch signals have mostly been reported in the context of episodic memory-like paradigms, where individual stimuli are only repeated a few times, whereas studies revealing prediction signals have generally involved a longer training phase to fully establish predictive associations before measuring neural signals. That is, when stimuli or associations are novel, the hippocampus is mainly driven by sensory signals that provide the opportunity to update our model of the world, i.e., prediction errors^28,38–41^. However, once learning is complete and environmental contingencies are no longer novel, hippocampal processing is dominated by retrieving predicted stimuli based on contextual cues to optimally guide perception^1,27,42,43^.

In line with this idea, recent work has shown that novel prediction errors can put the human hippocampus in an encoding mode^44^, increasing sensory processing (i.e., EC to CA1 connectivity) and decreasing mnemonic retrieval (CA3/DG to CA1 connectivity). Additionally, behavioural evidence suggests that expectation violation^45^ and novelty^46^ can bias the hippocampus towards performing pattern separation, proposed to underlie prediction error computations^28^. Indeed, in the context of episodic memory, it has been proposed that the hippocampus operates in two distinct modes, namely an encoding mode that prioritises processing of novel sensory signals and promotes plasticity, and a retrieval mode that priorities memory retrieval through pattern completion^47,48^. The hippocampus is thought to be biased towards the encoding mode by novelty-induced increases in neuromodulators such as acetylcholine (ACh) and norepinephrine (NE)^49–51^ and hippocampal theta phase resets^52–54^.

However, a proper test of this proposal requires establishing whether the hippocampus switches from representing errors to predictions as learning progresses. Here, we present the results of two high-resolution fMRI studies (N=24 for both experiments) directly testing this hypothesis. Participants were exposed to auditory cues that predicted the identity of an upcoming visual shape (with 75% validity). To preface our findings, we found that the hippocampus initially preferentially represented unexpected shapes, but later switched to representing the cue-predicted shape regardless of whether it was actually presented. Furthermore, in this latter phase we observed increased informational connectivity between the posterior subiculum and early visual cortex (V1), in line with hippocampal predictions being relayed to sensory cortex. These findings demonstrate a switch in hippocampal representations from errors to predictions as associative learning proceeds.

## Results

We present the results of two human fMRI studies (N=24 participants in both experiments) in which human participants were exposed to auditory cues that predicted the identity of an upcoming visual shape (with 75% validity) (Figure 1A-B). On each block of trials (n=32 trials per block in Experiment 1 and n=128 in Experiment 2) new auditory cues were presented, such that novel associations would have to be learnt.

**Figure 1.**
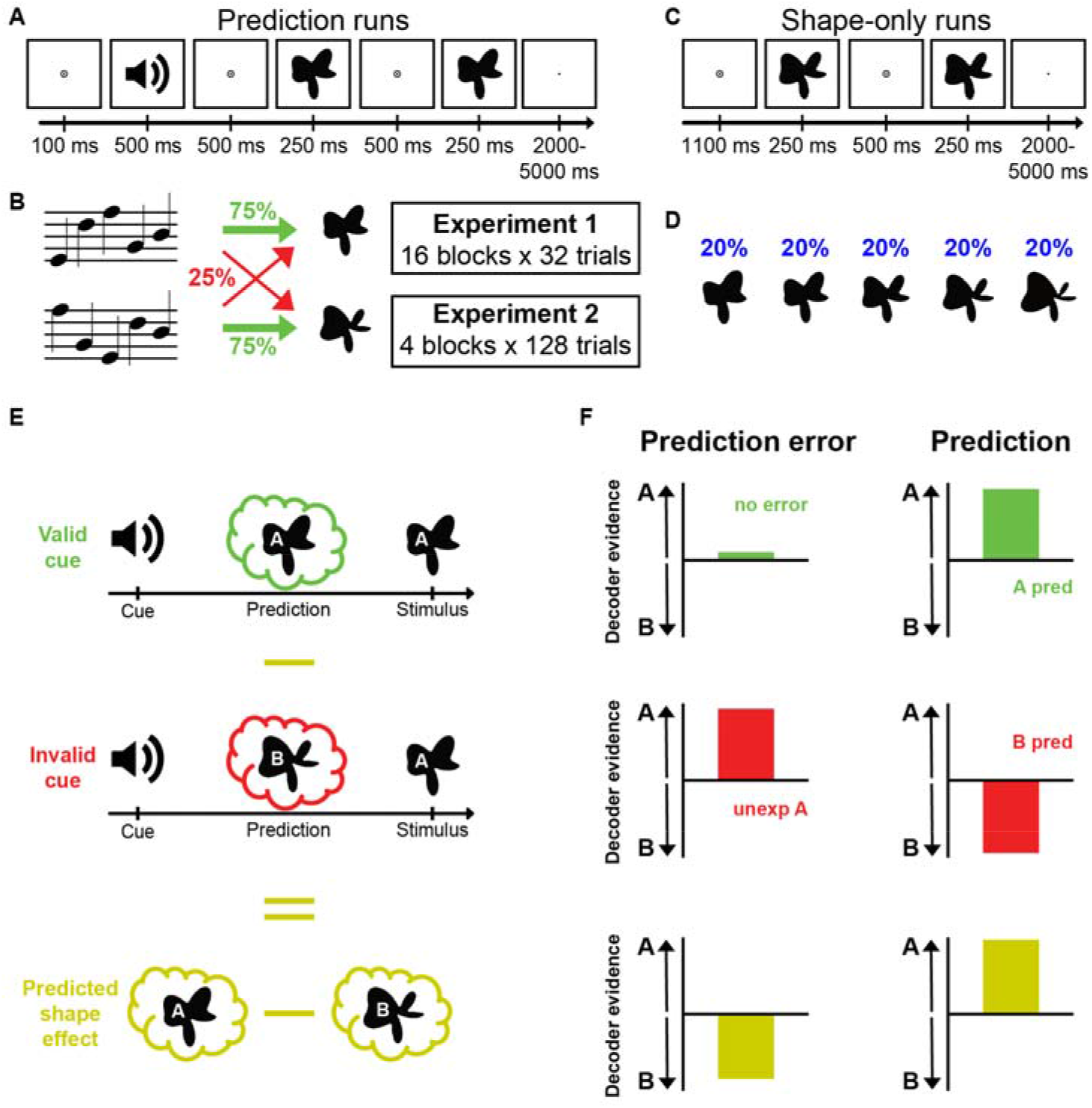
Experimental paradigm and analysis. **A)** During prediction runs, an auditory cue preceded the presentation of two consecutive shape stimuli. On each trial, the second shape was either identical to the first or slightly warped with respect to the first along an orthogonal dimension, and participants’ task was to report whether the two shapes were the same or different. **B)** The auditory cues predicted whether the first shape on a given trial would be shape 2 or shape 4 (of 5 shapes). The cue was valid on 75% of trials, whereas in the other 25% of (invalid) trials the unpredicted shape was presented. **C)** During shape-only runs, no auditory cues were presented. As in the prediction runs, two shapes were presented on each trial, and participants’ task was to report same or different. **D)** All five shapes appeared with equal (20%) likelihood during shape-only runs. **E)** Subtracting the response evoked by invalidly from validly predicted shapes isolated the effect of the predictive cues. **F)** Hypothesised shape decoding results if the hippocampus represents either prediction errors (left column) or predictions (right column).

Participants performed a shape discrimination task that was orthogonal to the predictive cues. Specifically, on each trial the first shape (validly predicted, 75% of trials, or invalidly predicted, 25%) was followed by a second shape that was either identical to the first (50% of trials) or very slightly warped (50%; see Methods for details). Participants’ task was to indicate whether the two shapes were the same or different. This task was designed to encourage participants to pay attention to the shapes while keeping the cue-shape contingencies task-irrelevant. In fact, participants were not informed that the auditory cues predicted the identity of the upcoming shape, and debriefing revealed that they did not become aware of this during the experiments. In other words, any learning of cue-shape associations was incidental and implicit.

Our experimental design allowed us to quantify how visual shape representations were modulated by the predictive auditory cues by subtracting signals evoked by validly and invalidly predicted shapes (Figure 1E-F). Multivoxel decoding analyses (Figure S1), trained on data from separate shape-only runs in which no predictive cues were presented (Figure 1C-D), were used to test whether hippocampal shape representations on valid and invalid trials (Figure 1E) were in line with either prediction error or prediction signals (Figure 1F). Specifically, if the hippocampus represents prediction error, one would expect it to represent unpredicted, but not predicted shapes (Figure 1F, left column). Alternatively, if the hippocampus represents predictions, it should represent the shape predicted by the auditory cue, regardless of which shape was actually presented (Figure 1F, right column)^27^.

### Experiment 1: short blocks

In Experiment 1, participants completed 16 blocks of 32 trials, with two novel auditory cues being presented in each block, while the same two visual shapes were presented throughout. Each auditory cue predicted which of the two shapes would be presented with 75% validity (Figure 1B).

#### Behavioural results

Participants were able to detect small differences in the shapes, during both the shape-only runs (67.7 +/- 1.7% correct; 29.7 +/- 1.8% modulation of the 3.18 Hz radial frequency component, mean +/- SEM) and during the prediction runs (69.0 +/- 1.4% correct; 28.7 +/- 1.9% modulation). Accuracy and reaction times (RTs) did not differ significantly between valid (68.9 +/- 1.5% correct; RT = 592 +/- 19 ms) and invalid (69.3 +/- 1.6% correct; RT = 595 +/- 20 ms; both *p* > 0.10) trials. This is as expected, since the discrimination task was orthogonal to the prediction manipulation (see Methods for details), and in line with previous results^27^.

#### fMRI decoding results

In the second half of the blocks, hippocampal activity patterns started to reflect unexpected (i.e. invalidly predicted) visual shapes (significant cluster from trial 22 to 32, *p* = 0.024; Figure 2A, red line). However, there was no significant representation of validly predicted shapes (no clusters with *p* < 0.05; Figure 2A, green line). In fact, there was a significant difference between invalidly and validly predicted shape decoding in the hippocampus (valid – invalid, significant cluster from trial 24 to 32, *p* = 0.028; Figure 2B). In other words, hippocampal activity patterns reflected shapes that were unexpectedly presented (e.g., decoding evidence for shape A when shape B was predicted but A was presented) but not shapes that were presented as expected (e.g., no evidence for shape A when shape A was predicted and presented). In sum, towards the end of the blocks, activity patterns in the hippocampus reflected a prediction error-like signal, representing unexpected but not expected shapes.

**Figure 2.**
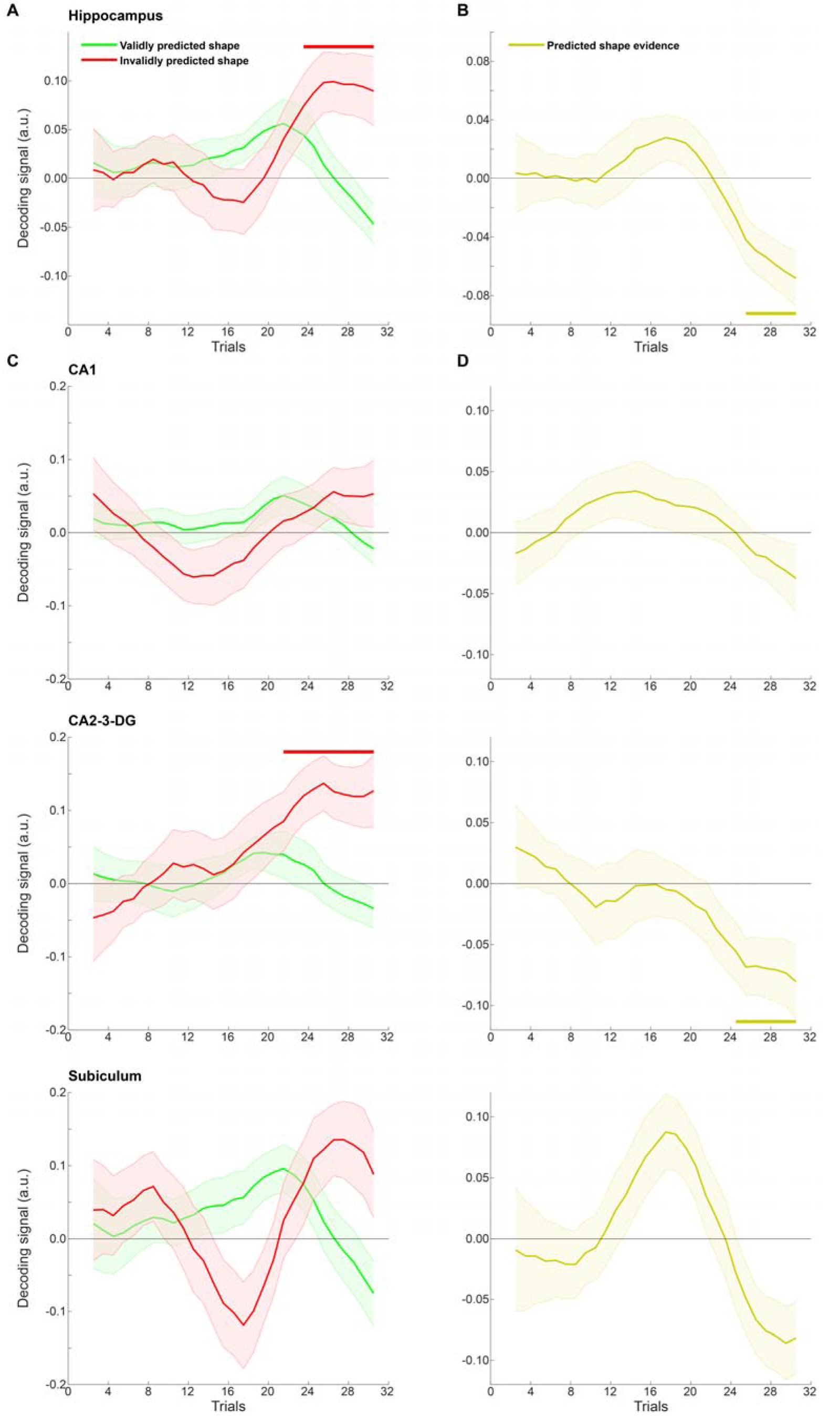
Experiment 1 shape decoding over trials. **A)** Decoding evidence for validly (green) and invalidly (red) predicted shapes in hippocampus. **B)** Decoding evidence for predicted (valid – invalid) shapes in hippocampus. **C)** Decoding evidence for validly (green) and invalidly (red) predicted shapes in hippocampal subfields. **D)** Decoding evidence for predicted (valid – invalid) shapes in hippocampal subfields. Horizontal lines indicate significant clusters. Shaded regions indicate SEM.

Segmenting the hippocampus into its subfields revealed that this effect was significantly present in CA2-3-DG (significant cluster for invalidly predicted shapes from trial 20 to 32, *p* = 0.014; significant cluster for valid - invalid from 23 to 32, *p* = 0.045), but not in CA1 and the subiculum (no clusters with *p* < 0.05), suggesting that this effect may have been driven by CA2-3-DG. However, decoding evidence for the predicted shape (i.e. valid – invalid, Figure 2D) in the last bin was not significantly different between the different subfields (*F*_2,46_ = 0.83, *p* = 0.44). Given recent interest in potential functional differences along the long axis of the hippocampus^55,56^, we also compared decoding evidence for the predicted shape in the last bin between the posterior and anterior hippocampus, but found no significant difference (*t*_23_ = 0.91, *p* = 0.37). However, decoding evidence for the predicted shape was significant in the posterior (*t*_23_ = -3.83, *p* = 0.00086) but not the anterior (*t*_23_ = -1.36, *p* = 0.19) hippocampus, suggesting the posterior hippocampus may be driving the prediction error-like effects.

In order to quantify the emergence of these signals over trials, we fit sigmoid functions, or S-curves, to the decoding evidence for predicted shapes in the hippocampus (Figure 3A; see Methods for details). In line with the results from the non-parametric cluster-based permutations tests reported earlier, the best fitting sigmoids had a significantly negative amplitude in the hippocampus (*t*_23_ = -2.17, *p* = 0.041) and CA2-3-DG (*t*_23_ = -2.90, *p* = 0.0080), but not in CA1 (*t*_23_ = -0.31, *p* = 0.76) and the subiculum (*t*_23_ = -1.38, *p* = 0.18). Finally, in a control analysis, to quantify the representational change over time without making any assumptions about the shape of this change, we calculated the derivative of the decoding evidence for the predicted shape over trials. In line with the previous analyses, in hippocampus (*t*_23_ = -2.72, *p* = 0.012) and CA2-3-DG (*t*_23_ = -2.84, *p* = 0.0092), but not in CA1 (*t*_23_ = -0.58, *p* = 0.57) and the subiculum (*t*_23_ = -1.55, *p* = 0.13), the average derivative over the course of the blocks was significantly negative.

**Figure 3.**
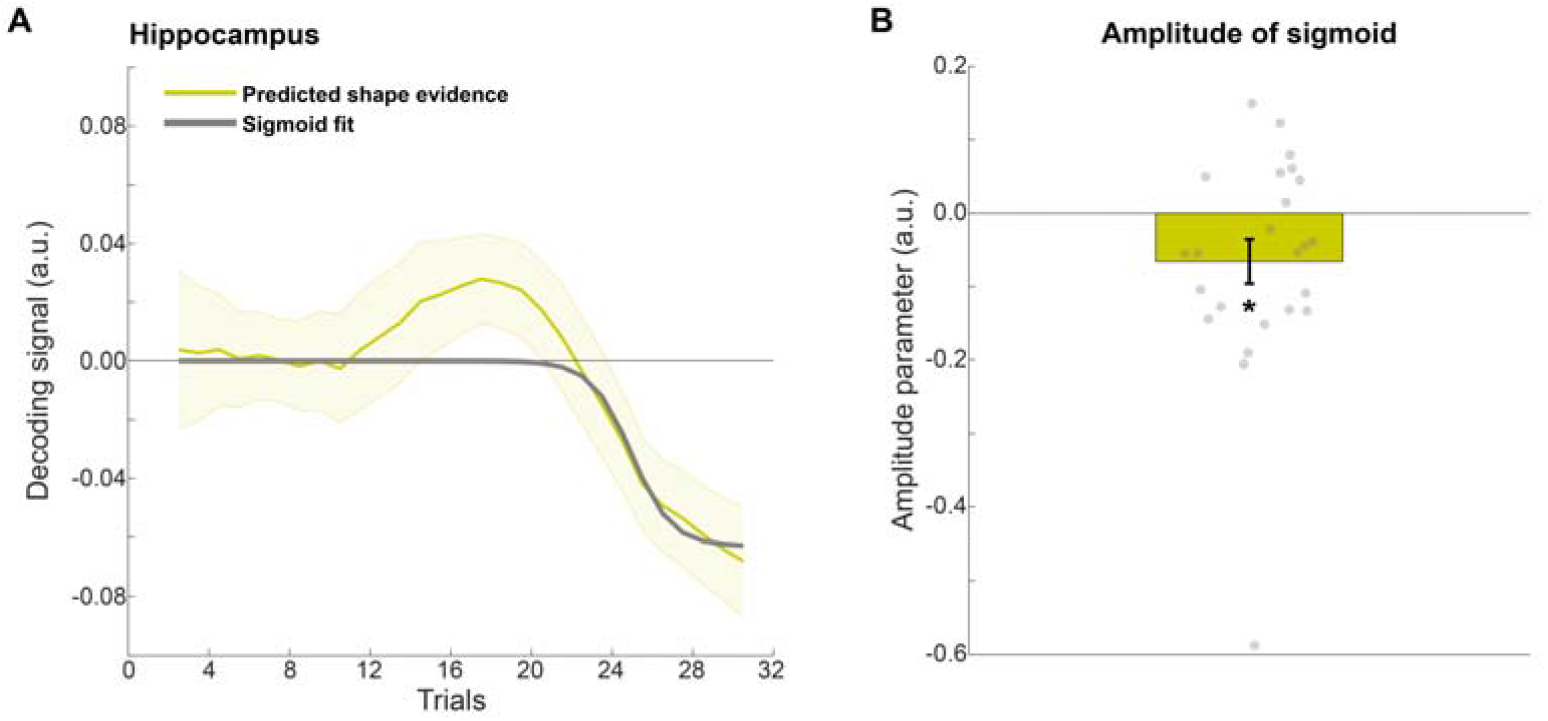
Quantification of hippocampal learning curve in Experiment 1. **A)** Sigmoid learning curve fit to predicted shape decoding in hippocampus. **B)** Amplitude parameter of sigmoid curve. Error bars indicate SEM. Dots indicate individual participants. *p < 0.05.

Based on previous findings of predictive signals in the caudate nucleus^4,27,57^, we also tested these effects in the caudate, and found that like the hippocampus, caudate activity patterns reflected unexpected (significant cluster from trial 20 to 29, *p* = 0.0062) but not expected (no clusters with *p* < 0.05) shapes towards the end of the blocks, with a significant difference between the two conditions (valid – invalid, significant cluster from trial 20 to 29, *p* = 0.033; Figure S2).

The fact that the hippocampus displayed a prediction error-like pattern (cf. Figure 2A and Figure 1F, left column) is striking given that several previous studies have reported prediction-like effects^5,26,37^. Specifically, a previous study with a virtually identical design^27^ revealed evidence for the shape predicted by the cue, regardless of which shape was actually presented (as in Figure 1F, right column). The crucial difference is that in these previous studies participants were exposed to the predictive associations for many trials before the fMRI session, whereas here participants learned novel predictive associations every block. Based on this, we hypothesised that the hippocampus may switch from representing prediction errors (early in learning) to representing predictions (once learning is complete) as learning progresses (Figure 4). In order to test this hypothesis, we performed a second fMRI experiment, in which participants (N=24) were exposed to the same cues for longer, and tested for potential switches in dynamics by fitting sigmoid learning curves to the decoding evidence over trials.

**Figure 4.**
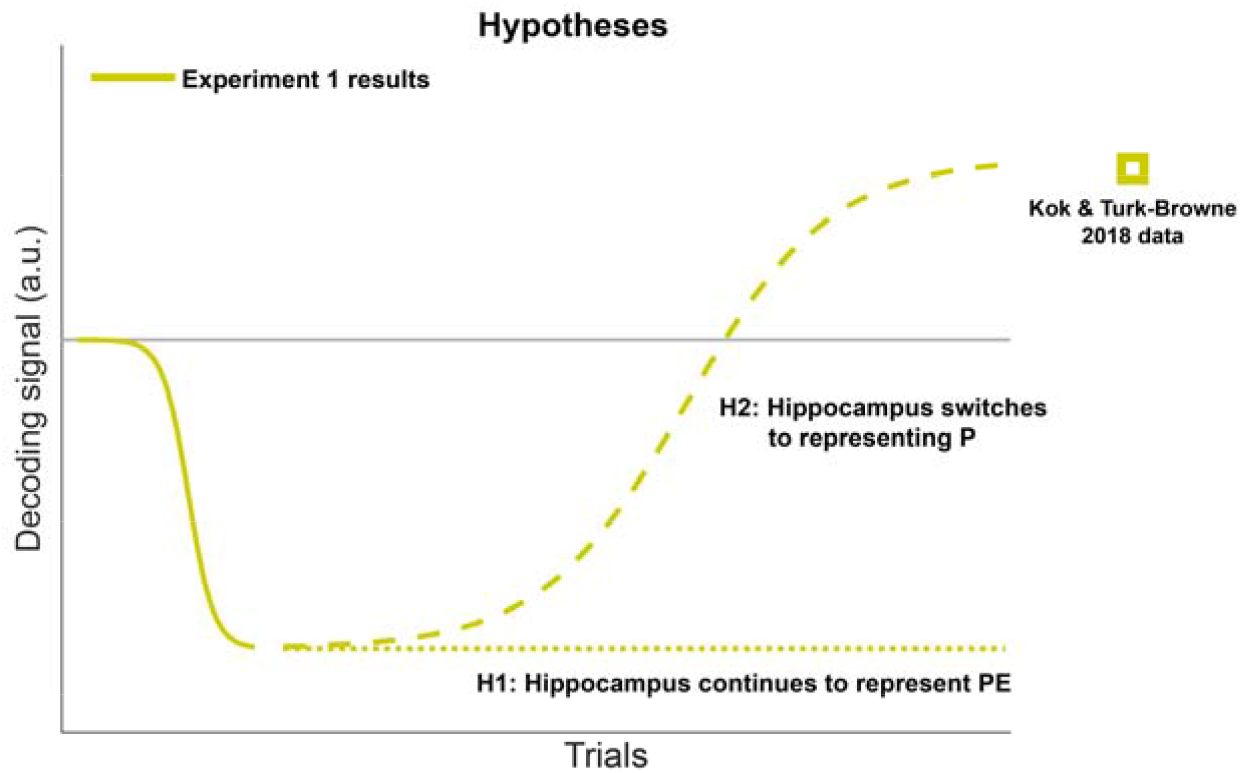
Hypotheses for Experiment 2. Experiment 1 revealed a negative shape decoding signal (solid line). Lengthening the learning phase may either result in this effect continuing (H1, dotted line), or lead to a switch towards positive prediction signals once learning is complete (H2, dashed line). Square indicates result from^27^, where participants were acquainted with the predictive cues before the fMRI session.

### Experiment 2: long blocks

In Experiment 2, participants were exposed to 4 blocks of 128 trials (compared to 16 blocks of 32 trials in Experiment 1), with new auditory predictive cues being presented in each block. In all other regards, Experiment 2 was identical to Experiment 1.

#### Behavioural results

As in Experiment 1, participants were able to detect small differences in the shapes, during both the shape-only runs (69.5 +/- 1.7% correct; 30.1 +/- 2.0% modulation of the 3.18 Hz radial frequency component, mean +/- SEM) and during the prediction runs (68.5 +/- 1.8% correct; 24.5 +/- 2.0% shape modulation). Accuracy and reaction times (RTs) again did not differ significantly between valid (68.6 +/- 1.9% correct; RT = 651 +/- 18 ms) and invalid (68.2 +/- 1.9% correct; RT = 654 +/- 18 ms; both *p* > 0.10) trials.

#### fMRI decoding results

Reflections of validly and invalidly predicted shapes in hippocampal activity patterns displayed striking of dynamics over time (Figure 5A). These dynamics were quantified by fitting two sigmoid curves to the decoding evidence for predicted shapes (Figure 5B), one with an inflection point in the first half of the blocks (trials 1-64) and the other with an inflection point in the second half (trials 65-128). This analysis revealed that an initial negative curve (amplitude parameter of early sigmoid; *t*_23_ = -2.26, *p* = 0.033), reflecting evidence for unexpected but not expected shapes (i.e., prediction error, as in Experiment 1) was followed by a positive curve (amplitude parameter of late sigmoid curve; *t*_23_ = 2.45, *p* = 0.022) about halfway through the blocks.

**Figure 5.**
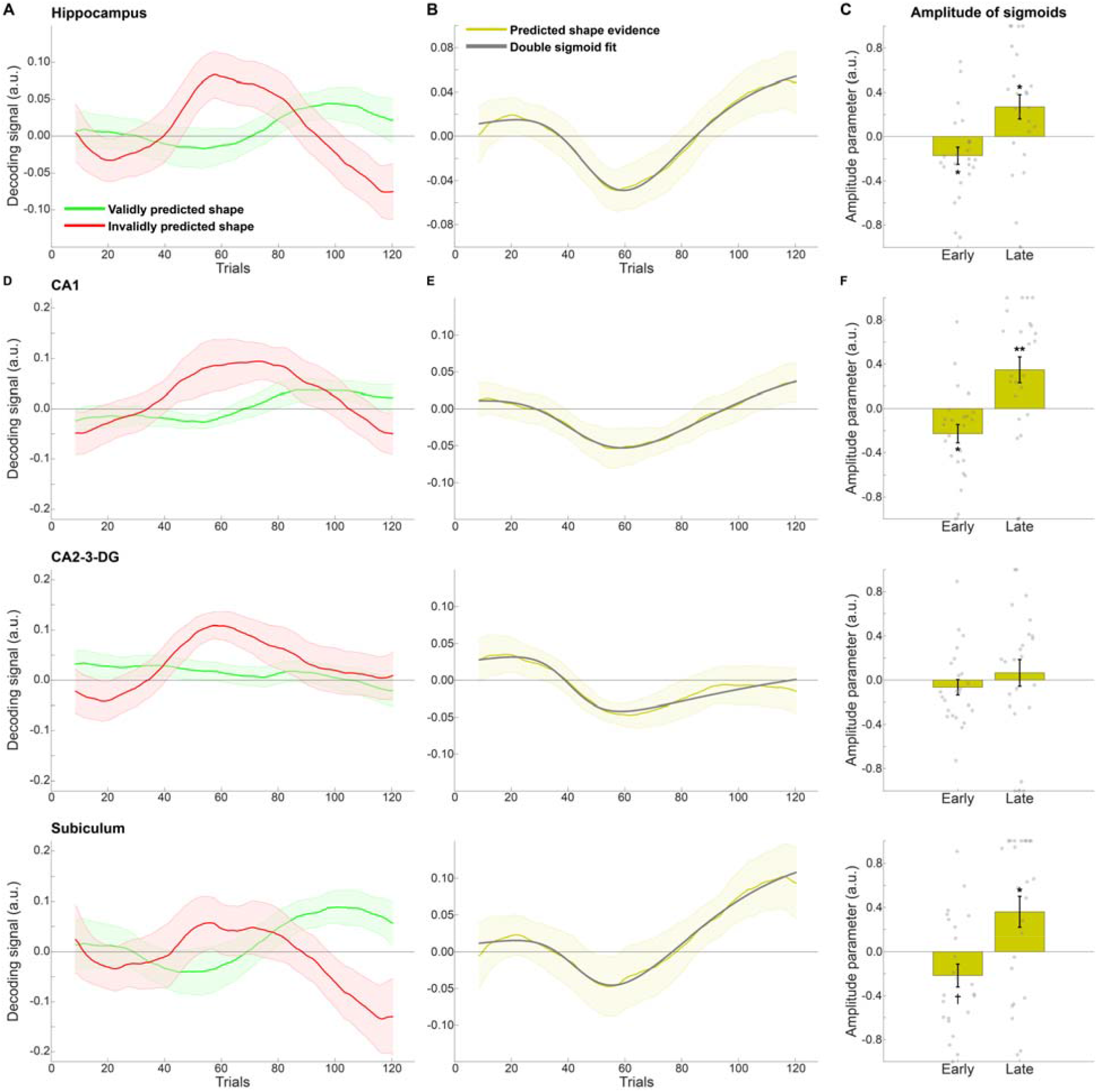
Experiment 2 shape decoding over trials. **A)** Decoding evidence for validly (green) and invalidly (red) predicted shapes in hippocampus. **B)** Decoding evidence for predicted (valid – invalid) shapes in hippocampus (yellow) with double sigmoid fit (gray). **C)** Amplitude parameters of early (midpoint between trials 1 and 64) and late (midpoint between trials 65 and 128) sigmoid curves in hippocampus. **D)** Decoding evidence for validly (green) and invalidly (red) predicted shapes in hippocampal subfields. **E)** Decoding evidence for predicted (valid – invalid) shapes in hippocampal subfields (yellow) with double sigmoid fit (gray). **F)** Amplitude parameters of early (midpoint between trials 1 and 64) and late (midpoint between trials 65 and 128) sigmoid curves in hippocampal subfields. Shaded regions and error bars indicate SEM. Dots indicate individual participants. *p < 0.05; **p < 0.01; † p = 0.05.

Segmenting the hippocampus into its subfields revealed that both the early negative and later positive learning curves were also significant in CA1 (early: *t*_23_ = -2.75, *p* = 0.011; late: *t*_23_ = 2.96, *p* = 0.0070), but not in CA2-3-DG (early: *t*_23_ = -0.95, *p* = 0.35; late: *t*_23_ = 0.54, *p* = 0.60), while in the subiculum the early negative curve was marginal (*t*_23_ = -2.07, *p* = 0.05) while the later positive one was significant (*t*_23_ = 2.56, *p* = 0.018).

As in Experiment 1, we performed a control analysis that did not make any assumptions about the shapes of the learning curves, in which we calculated the derivative of the decoding evidence for the predicted shape (Figures 5B and 5E), separately for the first and second half of the blocks. In line with the curve fitting results, the derivative was significantly different in the first versus the second halves of the blocks in hippocampus (*t*_23_ = -2.67, *p* = 0.014) and CA1 (*t*_23_ = -2.41, *p* = 0.024), while this difference was marginal in CA2-3-DG (*t*_23_ = -2.06, *p* = 0.051) and not significant in the subiculum (*t*_23_ = -1.34, *p* = 0.19). This was driven by the derivative being significantly positive in the second half of the blocks in hippocampus (*t*_23_ = 2.25, *p* = 0.034) and CA1 (*t*_23_ = 2.24, *p* = 0.035), but marginally negative in the first half (hippocampus: *t*_23_ = -2.00, *p* = 0.057; CA1: *t*_23_ = -1.98, *p* = 0.060). In CA2-3-DG, the derivative was significantly negative in the first half (*t*_23_ = -2.73, *p* = 0.012) but not the second half (*t*_23_ = 0.87, *p* = 0.39), while neither half was significant in the subiculum (first half: *t*_23_ = -0.51, *p* = 0.61; second half: *t*_23_ = 1.49, *p* = 0.15). There was no significant difference between the hippocampal subfields in terms of the derivative of the decoding timecourses in either the first (*F*_2,46_ = 0.58, *p* = 0.56) or second halves (*F*_2,46_ = 0.79, *p* = 0.46) of the blocks.

However, there was a significant difference between posterior and anterior hippocampus, with the positive derivative in the second half of the blocks being stronger in posterior than anterior hippocampus (*t*_23_ = 3.00, *p* = 0.0064; Figure S3). In fact, the early negative (posterior: *t*_23_ = -2.43, *p* = 0.024; anterior: *t*_23_ = -0.86, *p* = 0.40) and late positive (posterior: *t*_23_ = 2.70, *p* = 0.013; anterior: *t*_23_ = 0.98, *p* = 0.34) sigmoids, as well as the difference in the derivative between the first and second halve of the blocks (posterior: *t*_23_ = 2.74, *p* = 0.012; anterior: *t*_23_ = 1.42, *p* = 0.17), were significant in the posterior, but not anterior hippocampus. Decoding evidence for the predicted shape at the end of the blocks (i.e., in the final bin) was also stronger in the posterior (*t*_23_ = 2.15, *p* = 0.042) than the anterior (*t*_23_ = 0.31, *p* = 0.76; posterior vs. anterior: *t*_23_ = 2.14, *p* = 0.043) hippocampus.

As in Experiment 1, there was no difference in decoding evidence for the predicted shape at the end of the blocks between the hippocampal subfields (*F*_2,46_ = 2.61, *p* = 0.084; Figure 5E), but it is worth noting that the evidence for the predicted shape was numerically largest in the posterior subiculum (0.13; *t*_23_ = 2.55, *p* = 0.018; Figure S4), in line with prediction signals being significantly largest in the subiculum in previous work^27^. Since the subiculum is a major output relay from the hippocampus to neocortex^58,59^, we speculate that this may be in line with hippocampal prediction signals being communicated to sensory cortex, in order to guide perception. If this were the case, one would expect functional connectivity between hippocampus and neocortex to increase as predictive associations are established^60,61^. We tested this hypothesis in an exploratory analysis of informational connectivity between the posterior subiculum and entorhinal cortex (EC; a major interface between hippocampus and cortex) as well as visual cortex (V1, V2 and LO) (see Methods for details). This analysis revealed increased informational connectivity at the end of the blocks (final bin) versus at the start of the blocks (first bin) between the posterior subiculum and EC (*t*_23_ = 2.40, *p* = 0.025; Figure 6) and V1 (*t*_23_ = 2.88, *p* = 0.0084), but not V2 (*t*_23_ = 1.73, *p* = 0.097) and LO (*t*_23_ = -0.28, *p* = 0.78).

**Figure 6.**
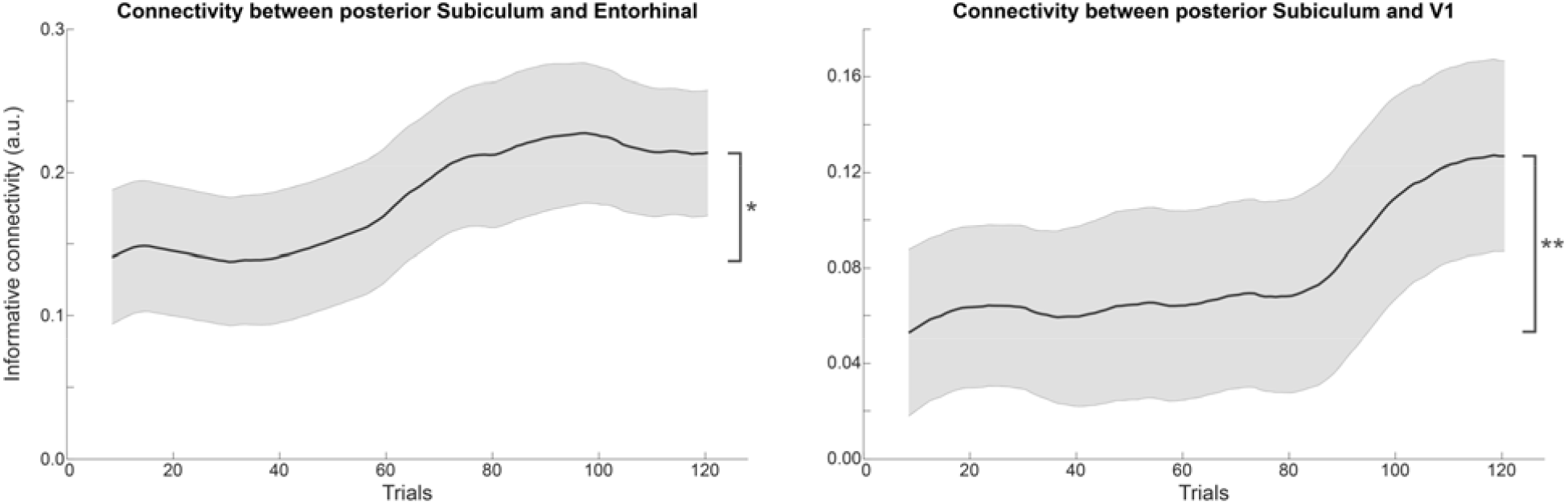
Experiment 2 informational connectivity between posterior subiculum and neocortex over trials. Time-resolved Pearson correlation between shape evidence in the posterior subiculum and entorhinal cortex (left panel) and V1 (right panel). Shaded indicate SEM. *p < 0.05; **p < 0.01.

The caudate nucleus displayed a qualitatively similar pattern of results as the hippocampus, with an initial negative sigmoid curve (*t*_23_ = -3.11, *p* = 0.0049) being followed by a positive curve (*t*_23_ = 3.65, *p* = 0.0013; Figure S5). The control analysis also revealed a significant difference in derivative in the first versus second half of the blocks (*t*_23_ = 2.90, *p* = 0.0081), driven by positive derivative in the second half (*t*_23_ = 3.94, *p* = 0.00066) but not the first half (*t*_23_ = -0.33, *p* = 0.74), and a significantly positive representation of the predicted shape in the final bin of the blocks (*t*_23_ = 3.06, *p* = 0.0055).

## Discussion

In two human fMRI experiments, we find that as learning of associative predictions progresses, the hippocampus switches from preferentially representing unexpected stimuli (i.e., prediction errors) to representing predicted shapes. These findings demonstrate a role for the hippocampus in both acquiring and exploiting predictive associations, and a switch between these modes depending on whether learning is ongoing (i.e., when prediction errors are informative^28^) or complete (only expected uncertainty remains). Concretely, what this suggests in the context of the current study is that prediction errors caused by early cue violations, when learning is still very much ongoing, dominate processing in the hippocampus, leading to the representation of the unexpectedly presented shape. On the other hand, once the 75%-25% cue contingencies are firmly learnt, the 25% cue violations are no longer treated as model updating (‘newsworthy’) events^28^, are therefore no longer upweighted, and the retrieval of the cued shape dominates. Note that we are not suggesting that this is an all-or-nothing switch; it is likely that the hippocampus always represents both predictions (through pattern completion in CA3^21,24,62,63^) and errors (potentially through mismatch comparison in CA1^33,35,64^), but that the balance between the two depends on contextual factors such as novelty and unexpected uncertainty. An analogous switch between prediction vs. surprise dominated representations has recently been proposed in the realm of perception, albeit on a sub-second time-scale^65^.

The early bias towards prediction errors is in line with recent demonstrations of hippocampal mode switches induced by novel prediction errors in humans^44^. Mechanistically, this switch may occur since novelty leads to an increase of neuromodulators like ACh and NE^49–51^, which suppress retrieval-related connections (CA3’s autorecurrence and CA3 -> CA1) relative to encoding-related ones (EC -> CA1)^66–68^. Alternatively, novelty may promote encoding on a faster time-scale by inducing a hippocampal theta phase reset^48,52–54^. Further research is needed to determine whether the switch demonstrated here was indeed driven by hippocampal mode changes or by a different mechanism that upweights novel prediction errors, such as attention^69–71^. For instance, methods with higher temporal resolution such as EEG/MEG or invasive electrophysiology could be used to investigate whether there is a relationship between hippocampal theta phase and error vs. prediction representations in the hippocampus. In either case, as learning progresses and novelty diminishes, a bias towards encoding prediction errors is abolished and retrieval of predictive associations dominates.

As prediction signals emerged in the hippocampus, functional connectivity increased between the posterior subiculum and the entorhinal cortex and primary visual cortex, demonstrating a potential route for relaying predictions to sensory cortex^26,60,72,73^. This relaying of predictions likely involves the same mechanisms that are responsible for hippocampus-mediated cortical reinstatement of memories^74–77^. Of course, fMRI connectivity analyses cannot determine directionality given the slow nature of the BOLD signal, so future research using electrophysiology^78,79^ or layer-specific fMRI^73,80^ will be required to test this hypothesis further.

It is noteworthy that the predictive associations studied here were fully implicit. Participants were not informed that there were any such associations, the predictions were incidental to the task, and debriefing indicated that participants did not become aware of them over the course of the experiments^8,43^. The fact that such implicit associations still involved the hippocampus is in line with theories of hippocampal processing based on the types of computations required, rather than whether they are explicit or implicit^23,39,81^. In fact, it has even been suggested that the hippocampus may engage in error-driven conjunction learning specifically when associations are incidental to the task subjects perform^82^.

In the current study, both prediction error and prediction signals seem to have been driven by the posterior rather than the anterior hippocampus. This finding is in line with suggestions that hippocampal representations increase in complexity and scale along the long axis^55^; simple cue-stimulus associations as studied here may therefore be encoded in posterior hippocampus^56^, whereas more complex representations such as narratives^55^ and scenes^83–85^ are encoded in anterior hippocampus.

Analysis of the caudate nucleus revealed similar prediction signals as in the hippocampus, in line with previous work employing a highly similar experimental design^27^, as well as other studies revealing involvement of the caudate in predictive processing^4,57^. Recently, it has been suggested that perceptual expectation signals in the tail of the striatum play a role in generating hallucination-like percepts in mice^86^. Future research is needed to establish whether the caudate and hippocampus play different or complementary roles in the processing of predictive associations^87,88^.

In the current study, novel predictive cues were introduced on each block of the experiment. It is an open question whether similar hippocampal dynamics would occur if the cue identities remained the same throughout the experiment, but the predictive contingencies switched. In other words, does the hippocampal switch observed here depend on the cues themselves being novel, or is it sufficient for only their predictive values to change, i.e., for there to be unexpected uncertainty^51^?

Additionally, whether the hippocampus signals predictions or prediction errors may also depend on the type of predicted stimulus. For instance, in previous work we reported hippocampal prediction signals for complex shapes, but prediction error-like signals for low-level features, i.e., the predicted orientation of a grating stimulus^89^. Future work systematically manipulating the complexity of visual stimuli may shed light on this by exploring the relationship between hippocampal computations and stimulus complexity^90^.

In sum, the current findings demonstrate a role for the hippocampus in both acquiring and exploiting predictive associations, bridging the fields of learning and perception. These fields have separately made progress in investigating the roles of prediction, novelty and uncertainty^1,51^, but have until now largely remained segregated literatures, despite great promise to inform one another^65,91^. Ultimately, weighting predictions and errors according to their reliability is crucial to optimally perceiving and engaging with our environment, and the current findings suggest that the hippocampus plays a crucial role in this process.

## Methods

### Participants

Both experiments aimed to recruit 24 healthy, right-handed, MR-compatible participants with normal or corrected-to-normal vision. All participants provided informed consent through a protocol reviewed by the University College London (UCL) Research Ethics Committee and were compensated a total of £27.50 for their time. Twenty-nine individuals completed Experiment 1, of which five were excluded due to our strict head motion criteria (five or more movements larger than 1.5 mm in any direction between successive functional volumes). The final sample consisted of 24 participants (12 female; age 25.6 ± 7.2, mean ± SD). Twenty-nine individuals completed Experiment 2, of which two were excluded for not performing the task above chance, and three due to excessive head motion (see criteria above). The final sample consisted of 24 participants (19 female; age 26.2 ± 7.0, mean ± SD).

### Stimuli

Visual and auditory stimuli were generated using MATLAB (Mathworks, Natick, MA, USA) and the Psychophysics Toolbox^92^. In the MR scanner, visual stimuli were displayed on a rear projection screen using a projector (1600 × 1200 resolution, 60 Hz refresh rate) against a grey background. Participants viewed the visual display through a mirror that was mounted on the head coil. The visual stimuli consisted of complex shapes defined radial frequency components (RFCs)^93,94^, identical to the shapes used in Kok & Turk-Browne^27^ (Figure 1). The contours of the stimuli were defined by seven RFCs, and a one-dimensional shape space was created by varying the amplitude of three out of the seven RFCs^27^. Specifically, the amplitudes of the 1.11 Hz, 1.54 Hz and 4.94 Hz components increased together, ranging from 0 to 36 (first two components), and from 15.58 to 33.58 (third component). Note that we chose to vary three RFCs simultaneously, rather than one, to increase the perceptual (and neural) discriminability of the shapes. Five shapes (Figure 1D) were selected from this continuum such that they represented a perceptually symmetrical sample of this shape space (see Kok & Turk-Browne^27^ for details). Additionally, a fourth RFC (the 3.18 Hz component) was used to create slightly warped versions of the five shapes, to enable the same/different shape discrimination cover task (see below). Experiments 1 and 2 presented identical shapes (black, subtending 4.5°), centred on fixation.

In the scanner, auditory stimuli were presented using MR-compatible ear buds (E-A-RTONE 3A, 10 Ohm, Etymotic Research, Elk Grove Village, IL, USA). The auditory stimuli consisted of sequences of pure tones, ranging in frequency from 261.36 Hz (C_4_) to 987.77 (B_5_) Hz (set of 14 tones: C_4_, D_4_, E_4_, F_4_, G_4_, A_4_, B_4_, C_5_, D_5_, E_5_, F_5_, G_5_, A_5_, B_5_; duration = 100 ms; 10 ms linear rise and fall ramps). Seventeen sequences of five tones (500 ms) were created by selecting the least correlated sequences from all permutations of {1, 2, 3, 4, 5}. These 17 sequences (e.g., 1-2-3-4-5, 1-5-4-3-2, 3-1-5-4-2, etc.) were further differentiated by assigning them different starting tones, in steps of 3. For instance, if sequence 1 was 1-2-3-4-5, sequence 2 was 4-8-7-6-5, sequence 3 was 9-7-11-10-8, etc. Since the maximum starting tone was 10, given the set of 14 tones, every fifth sequence started with starting tone 1 again. For each sequence, a mirrored sequence was generated in order to created 17 pairs of easily distinguishable sequences consisting of the same tones (e.g., sequence 1-2-3-4-5 was paired with 5-4-3-2-1, sequence 4-8-7-6-5 was paired with 8-4-5-6-7, etc.). In Experiment 1, 16 of these pairs were assigned in random order to the sixteen blocks, while the seventeenth pair was used during the practice block outside the scanner (see below). In Experiment 2, only the first four pairs were used, since this experiment contained only four blocks (see below), while the fifth pair was used for practice.

### Experimental procedure

The trial structure was identical in both experiments. The start of each trial was signalled by the presentation of a fixation bullseye (diameter, 0.7°). During prediction runs, an auditory cue (sequence of five tones; 500 ms) was presented 100 ms after the trial onset (Figure 1A). Following a 500 ms delay, two consecutive shapes were presented for 250 ms each, separated by a 500ms fixation screen. The auditory cues predicted whether the first shape on that trial would be shape 2 or shape 4 (out of five shapes; Figure 1B & 1D). The cue was valid on 75% of trials, whereas in the other 25% of trials the unpredicted shape would be presented. For instance, a specific auditory cue might be followed by shape 2 on 75% of trials and by shape 4 on the remaining 25% of trials. On each trial, the second shape either was identical to the first (50%), or slightly warped (50%), by modulating the amplitude of the 3.18 Hz RFC component defining the shape. This modulation could be either positive or negative (counterbalanced over conditions) and participants’ task was to indicate whether the two shapes on a given trial were the same or different, using an MR-compatible button box (750 ms response interval). This task was designed to encourage participants to attend the visual shapes, while avoiding a relationship between the perceptual prediction and the task response. Furthermore, by modulating one of the RFCs that was not used to define our one-dimensional shape space, we ensured that the shape change on which the task was performed was orthogonal to the changes that defined the shape space, and thus orthogonal to the shape features predicted by the auditory cues. The size of the shape modulation was determined by a staircasing procedure^95^, updated after every trial to ensure sufficient task difficulty (∼75% correct). The end of each trial was signalled by replacing the fixation bullseye with a single fixation dot, encouraging participants to continue to fixate (inter trial interval jittered between 1.25 and 4.25 s).

Experiment 1 consisted of 16 blocks of 32 trials, presented in four prediction runs (4 blocks per run, 30s breaks between runs, ∼12 min per run). In each block, a different pair of cues was presented. For each trial number (1 to 32) we counterbalanced 1) which cue was presented, 2) whether the cue was valid (75%) or invalid (25%), and 3) whether the two shapes were the same or different.

Experiment 2 consisted of 4 blocks of 128 trials (1 block per prediction run, 30s break halfway, ∼12 min per run), with a different pair of cues presented in each block. As in Experiment 1, cue validity was counterbalanced for every trial position, but given the smaller number of blocks, the presented cue and shape modulation were counterbalanced over groups of four trial positions (trials 1-4, 5-8, etc.) rather than for every trial position. This was reflected in the analyses by a four-fold increase in the trial averaging window; see below.

In both experiments, which pair of cues was assigned to which block, as well as which member of each pair predicted which shape, was counterbalanced across participants.

In addition to the four prediction runs, both experiments also contained two shape-only runs, flanking the prediction runs, constituting the first and last (sixth) runs of the experiments. In these runs (120 trials per run, ∼12 min) no auditory cues were presented (Figure 1C). As in the prediction runs, each trial started with the appearance of a fixation bullseye followed 1100ms later by two shapes (250 ms each, 500 ms interval). On each trial one of the five possible shapes was presented, with equal (20%) likelihood (Figure 1D). As in the prediction runs, participants’ task was to indicate whether the two shapes were the same or different. The size of the shape modulations was controlled by a staircase separate from that of the prediction runs, to equate task difficulty in these runs with five instead of two possible initial shapes. The shape-only runs acted as the training data for our shape decoding model, see below.

Before both experiments, participants completed an instruction and practice session to acquaint them with the task (∼30 min). During practice, participants completed 100 shape-only trials and 16 prediction trials. The pair of auditory cues used during the short prediction run was not included in the main experiments.

After the experiments, participants completed a short questionnaire that indicated whether or not they became aware of the predictive nature of the auditory cues. The responses to both an open-ended question (“Can you tell us what the meaning of the sounds was during the experiment?”) as well as a guided one (“During every block of the experiment, two different sounds were played. These sounds predicted which shapes would appear. For instance, a series of rising tones might predict that you’ll see shape A, and falling tones might predict you’ll see shape B. These predictions were 75% valid, so on 25% of trials they were incorrect. Did you realise this?”) indicated that the vast majority of participants did not become aware of the predictions in either experiment (Experiment 1: 1 out of 22 participants indicated that they realised the cues predicted which shape would appear, no data for 2 participants; Experiment 2: 0 out of 22 participants indicated that they realised the cues predicted which shape would appear, no data for 2 participants).

### MRI acquisition

In both experiments structural and functional MRI data were collected on a 3T Siemens Prisma scanner with a 64-channel head coil at the Wellcome Centre for Human Neuroimaging (WCHN). Note that in two different scanners with identical specifications were used for the two experiments, for availability reasons. Functional images for both experiments were acquired using a T2*-weighted multiband echo-planar imaging sequence (TR = 1000 ms; TE = 33.0 ms; 60 transverse slices; voxel size = 1.5 × 1.5 × 1.5 mm; flip angle = 55°, multiband factor = 6). This sequence produced a partial volume for each participant, which covered the occipital and temporal lobes, including and parallel to the hippocampus. Field map data were acquired using a Siemens Field Map sequence (TR = 1020.0ms; short TE = 10.00ms; long TE = 12.46ms; voxel size = 3.0 × 3.0 × 2.0 mm, 64 transverse slices, flip angle = 90°). Anatomical images were acquired using a T1-weighted Magnetisation Prepared Rapid Gradient Echo (MPRAGE), using a Generalized Auto calibrating Partially Parallel Acquisition (GRAPPA) factor of 2 (TR = 2530 ms; TE = 3.34ms; 176 sagittal slices; voxel size = 1.0 × 1.0 × 1.0 mm; flip angle = 7°). To enable hippocampal segmentation, a T2-weighted turbo spin-echo (TSE) image (TR = 12650 ms; TE = 45 ms; voxel size = 0.4 × 0.4 × 1.5 mm; 54 coronal slices perpendicular to the long axis of the hippocampus; flip angle = 122°) was acquired.

### fMRI preprocessing

Images for both experiments were preprocessed using Statistical Parametric Mapping (SPM12, http://www.fil.ion.ucl.ac.uk/spm, Wellcome Centre for Human Neuroimaging, London, UK). The first six volumes of each functional run were discarded to allow T1 equilibration. For each run, the remaining functional images were spatially realigned to correct for head motion, and simultaneously supplied to B0 unwarping, using SPM’s realign and unwarp function. The functional data were temporally high-pass filtered with a 128s period cut-off. No spatial smoothing was applied, and all analyses were performed in participants’ native space. The T1 and T2-weighted structural scans were co-registered and subsequently co-registered to the mean functional scan.

### Regions of Interest

The hippocampus and its subfields, CA1, CA2-3-DG, and the subiculum, were defined based on the structural T2 and T1 images using the automatic segmentation of hippocampal subfields (ASHS)^96^ machine learning toolbox, in conjunction with a database of manual medial temporal lobe (MTL) segmentations from a separate set of 51 participants^97,98^. Consistent with previous studies, CA2, CA3 and DG were combined into a single region of interest (ROI) since these subfields are difficult to distinguish at our functional resolution (1.5 mm isotropic). This method also yielded an entorhinal cortex (EC) ROI for our informative connectivity analysis (see below). Results of the automated segmentation were inspected visually for each participant. Additionally, a caudate region of interest (ROI), as well as visual cortex ROIs for our informational connectivity analysis – V1, V2 lateral occipital cortex (LO) – were automatically defined in each participant’s T1-weighted anatomical scan using FreeSurfer (http://surfer.nmr.mgh.harvard.edu/). The visual cortex ROIs were restricted to the 500 most active voxels during the shape-only runs, to ensure that we were measuring responses in the retinotopic locations corresponding to our visual stimuli. Since no clear retinotopic organization is present in the other ROIs, cross-validated feature selection was used instead (see below). All ROIs were collapsed over the left and right hemispheres, as we had no hypotheses regarding hemispheric differences.

### fMRI data modelling

For both experiments, the pattern of activity evoked by each single trial of the prediction runs, in each ROI, was estimated using the Least-Squares-Separate method^99,100^. That is, a separate GLM was created for every trial, such that each trial is modelled once as a regressor of interest, with all other trials combined into a single nuisance regressor. Delta functions were inserted at the onset of the trial of interest (first regressor) and all other trials (second regressor) and convolved with a double-gamma hemodynamic response function (HRF) and its temporal derivative^101^. The voxel-wise parameter estimates for the trial-of-interest HRF regressor constituted the estimated BOLD activity pattern for each trial. This method has been shown to improve the estimation of single-trial BOLD responses, compared with a GLM with one regressor for each trial^99^. In addition to these regressors, the GLMs included nuisance regressors consisting of the head motion parameters resulting from spatial realignment, their derivatives, and the square of these derivatives (i.e., 18 motion parameters in total). The data from the shape-only runs were analysed using a more conventional GLM, with one regressor for each of the five shapes and 18 head motion nuisance regressors.

### Shape decoding

In order to probe neural shape representations, a forward modelling approach was used to decode the shapes from the patterns of BOLD activity in each ROI^27,102^. The decoding algorithm was identical to that used in Kok & Turk-Browne^27^, and will be outlined here briefly (see Figure S1 for a visual depiction).

The shape selectivity of each voxel was characterised as a weighted sum of five hypothetical channels, each with an idealised shape tuning curve (or basis function), consisting of a halfwave-rectified sinusoid raised to the fifth power. In the first stage of the analysis, the parameter estimates obtained from the two shape-only runs were used to estimate the weights on the five hypothetical channels separately for each voxel, using linear regression. Specifically, let *k* be the number of channels, *m* the number of voxels, and *n* the number of measurements (i.e., the five shapes). The matrix of estimated response amplitudes for the different shapes during the shape-only runs (**B**_**train**_, *m x n*) was related to the matrix of hypothetical channel outputs (**C**_**train**_, *k x n*) by a weight matrix (**W**, *m x k*):

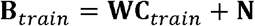

The weight matrix was estimated by least squares estimation:

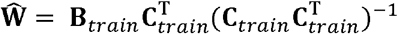

Using these weights, the second stage of analysis consisted of reconstructing the channel outputs associated with the pattern of activity across voxels evoked by each trial in the prediction runs (**B**_**test**_), again using linear regression:

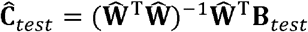

Where **Ĉ**_*test*_ are the estimated channel outputs. These channel outputs were used to compute a weighted average of the five basis functions, reflecting a neural shape tuning curve (Figure S1). Note that, during the main experiment (i.e., the prediction runs), only shapes 2 and 4 were presented. Decoding performance was quantified by subtracting the amplitude of the shape tuning curve at the presented shape (e.g., shape 2) from the amplitude at the non-presented shape (shape 4). This procedure yielded a measure of decoding evidence for the presented shape on each trial, in each ROI.

For all ROIs, voxel selection was based on data from the shape-only runs, in which no predictions were present, to ensure voxel selection was independent of the data in which we tested our effects of interest (i.e., the prediction runs). In visual cortex ROIs, we selected the 500 most active voxels during the shape-only runs. However, the hippocampus and caudate did not show a clear evoked response to visual stimuli, as defined by a lack of significant fit of a regressor of stimulus onset times convolved with a canonical haemodynamic response to the mean hippocampal time course. Therefore, we applied a different method of voxel selection for these ROIs. Voxels were first sorted by their informativeness, that is, how different the weights for the different channels were from each other, as indexed by the standard deviation of the weights. Second, the decoding model was trained and tested on different subsets of these voxels (between 10 and 100%, in 10% increments), within the shape-only runs (trained on one run and tested on the other). For all iterations, decoding performance on shapes 2 and 4 was quantified as described above, and the number of voxels that yielded the highest performance was selected. This procedure was used for voxel selection in the hippocampus (Experiment 1: 1068 voxels selected; Experiment 2: 970 voxels; group average), CA1 (Experiment 1: 271 voxels; Experiment 2: 313 voxels), CA2-3-DG (Experiment 1: 374 voxels; Experiment 2: 433 voxels), subiculum (Experiment 1: 273 voxels; Experiment 2: 249 voxels), and caudate (Experiment 1: 1292 voxels; Experiment 2: 1249 voxels).

### Quantifying time courses of shape representations

A sliding window approach was used to investigate how shape representations evolved over trials. In Experiment 1, this window consisted of 4 trial positions (i.e., trials 1-4 of all 16 blocks, followed by trials 2-5, trials 3-6, etc.), while for Experiment 2 the window was four times as wide (16 trial positions; trials 1-16 of all 4 blocks, trials 2-17, trials 3-18, etc.) to compensate for the four-fold decrease in the number of blocks (i.e. the number of trials-per-position). Within each window, we averaged the decoding evidence for validly and invalidly predicted shapes separately. In order to quantify evidence for the shape predicted by the cue, controlling for the actually presented shape, evidence for validly and invalidly predicted shapes was subtracted (i.e., averaging (1 - evidence) for the invalidly predicted shapes with evidence for the validly predicted shapes) (Figure 1E-F). Finally, the decoding time courses were smoothed by averaging over a sliding window. In Experiment 1 each bin was averaged with the previous and subsequent 4 bins, yielding a window size of 9 bins. In Experiment 2 the window size was 33 bins, containing the previous and subsequence 16 bins.

Initially, in Experiment 1, in a fully assumption-free analysis, we performed cluster-based permutation tests^103^ on the time courses, to test whether the decoding signals differed significantly from zero at any timepoint. Specifically, univariate *t* statistics were calculated for all timepoints, and neighbouring elements that passed a threshold value corresponding to a *p* value of 0.05 (two-tailed) were collected into clusters. Cluster-level test statistics consisted of the sum of *t* values within each cluster, which were compared to a null distribution created by drawing 10,000 random permutations of the observed data. A cluster was considered significant when its *p* value was below 0.05 (i.e., a cluster of its size occurred in fewer than 5% of the null distribution clusters).

Subsequently, the obtained time courses of decoding evidence for the predicted shapes were quantified by fitting sigmoid curves to them. In Experiment 1, this consisted of a single sigmoid:

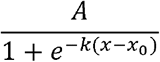

With a midpoint *x*_0_ between trial 1 and 32, slope *k* between 0.01 and 1, and amplitude A between - 1 and 1. These parameters were fitted using Matlab’s fmincon function, wrapped in GlobalSearch. We ran 100 iterations with random parameter starting values (within their prescribed ranges), in order to avoid local minima. The amplitude parameter was submitted to simple **t**-tests to test whether learning curves significantly deviating from zero. In Experiment 2, this consisted of a combination of two sigmoids, to test whether dynamics changed as learning progressed:

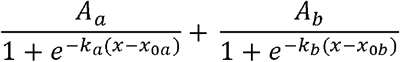

With slopes *k*_*a*_ and *k*_*b*_ between 0.01 and 1, amplitudes *A*_*a*_ and *A*_*b*_ between -1 and 1. The first sigmoid had a midpoint *x*_*0a*_ between trial 1 and 64, while the second had a midpoint *x*_*Ob*_ between trial 65 and 128, allowing them to capture potential differences between the first and second half of the blocks. Note that both sigmoids’ amplitudes were free to range between -1 and 1, meaning that this analysis contained no priors on the signs of the curves. As in Experiment 1, the amplitude parameters were submitted to simple **t**-tests to test whether learning curves significantly deviating from zero.

In a control analysis that made no assumptions on the shapes of the time courses, we calculated the average derivatives of the decoding time courses. For Experiment 2, this was done separately for the first (trials 1-64) and second (trials 65-128) half of the blocks, to be investigate whether dynamics changed as learning progressed.

All analyses were initially performed on the hippocampus ROI as a whole, and when significant this was followed up by investigating hippocampal subfields.

### Informational connectivity

In an exploratory analysis, we investigated whether functional connectivity between regions (specifically, between the posterior subiculum and EC, V1, V2, and LO) changed over trials in Experiment 2. Specifically, the Pearson correlation in decoding evidence over trials between two regions was calculated^104^, within the sliding windows described above. This analysis yielded time courses of correlation values, with a positive value indicating that whenever region A represents shape 2 (rather than shape 4), region B is likely to do so as well. Changes in informational connectivity over time were tested by comparing *r* values at the end of the blocks (i.e., the final window, containing trials 113-128) with the start of the blocks (the first window, containing trials 1-16), using paired-sample **t**-tests.

## Data availability statement

All region-specific fMRI time course data and corresponding analysis code will be made available on the OSF platform (DOI to follow).

## Acknowledgements

The authors would like to thank Patricia Andrea Cabiles, Victoire Martignac and Ellis Langford for assistance with data collection, and Anna Schapiro for helpful discussion of these findings. This work was supported by a Wellcome/Royal Society Sir Henry Dale Fellowship [218535/Z/19/Z] and a European Research Council (ERC) Starting Grant [948548] to P.K. The Wellcome Centre for Human Neuroimaging is supported by core funding from the Wellcome Trust [203147/Z/16/Z].

## Author contributions

P.K. designed the study; F.A. collected the data; F.A. and P.K. analysed the data and wrote the manuscript.

## Supplementary Figures

**Figure S1.**
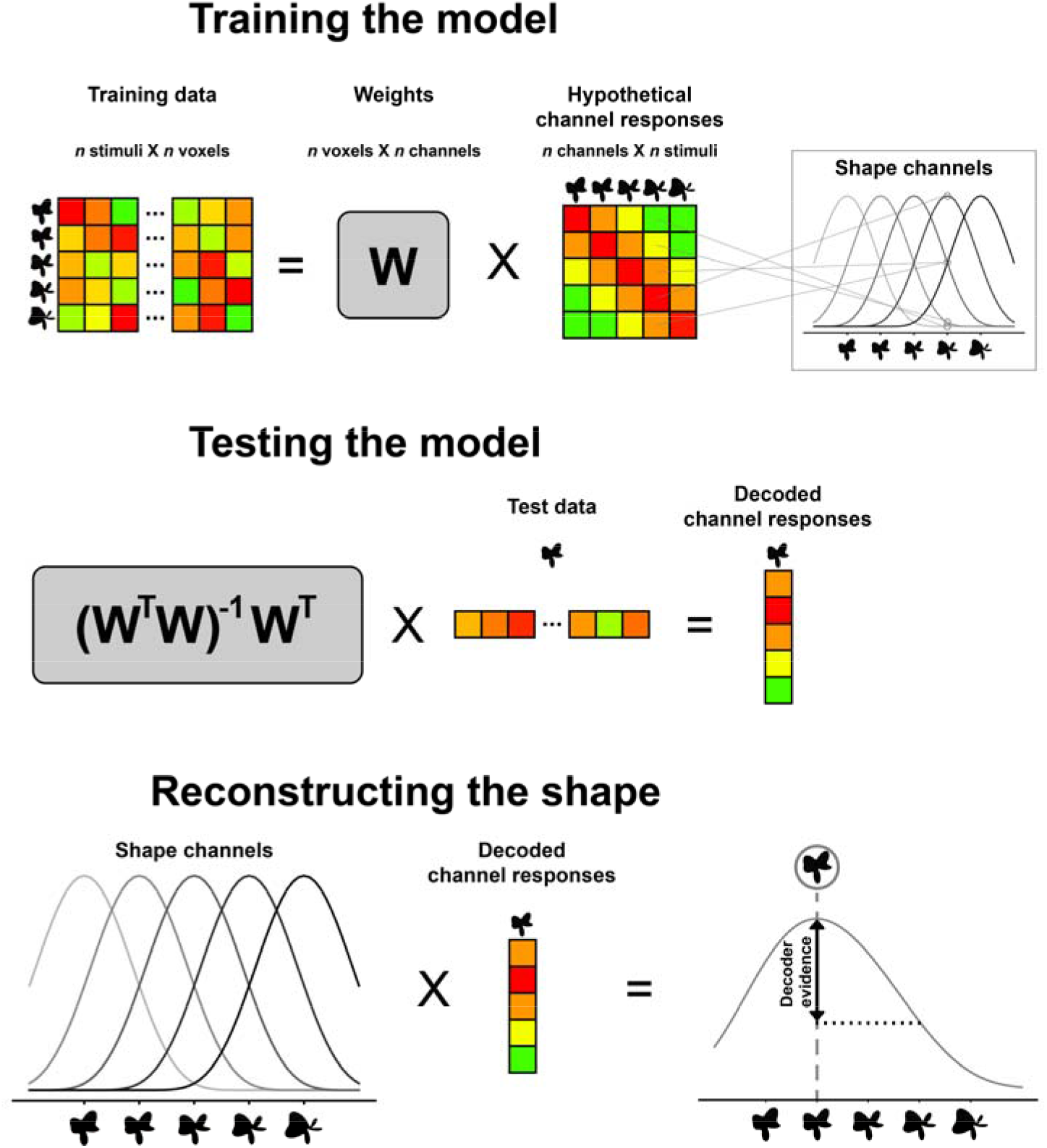
Illustration of shape decoding analysis. In order to probe neural shape representations, a forward modelling approach was used to decode the shapes from the patterns of BOLD activity in each ROI. In the first stage of the analysis, the parameter estimates obtained from the two shape-only runs were used to estimate the weights on the five hypothetical channels separately for each voxel, using linear regression. This constituted training the model. The second stage, testing the model, consisted of estimating the channel outputs associated with the pattern of activity across voxels evoked by each trial in the prediction runs, again using linear regression. These estimated channel outputs were used to compute a weighted average of the five basis functions, reflecting a neural shape tuning curve. Note that, during the main experiment (i.e., the prediction runs), only shapes 2 and 4 were presented. Decoding performance was quantified by subtracting the amplitude of the shape tuning curve at the presented shape (e.g., shape 2) from the amplitude at the non-presented shape (shape 4).

**Figure S2.**
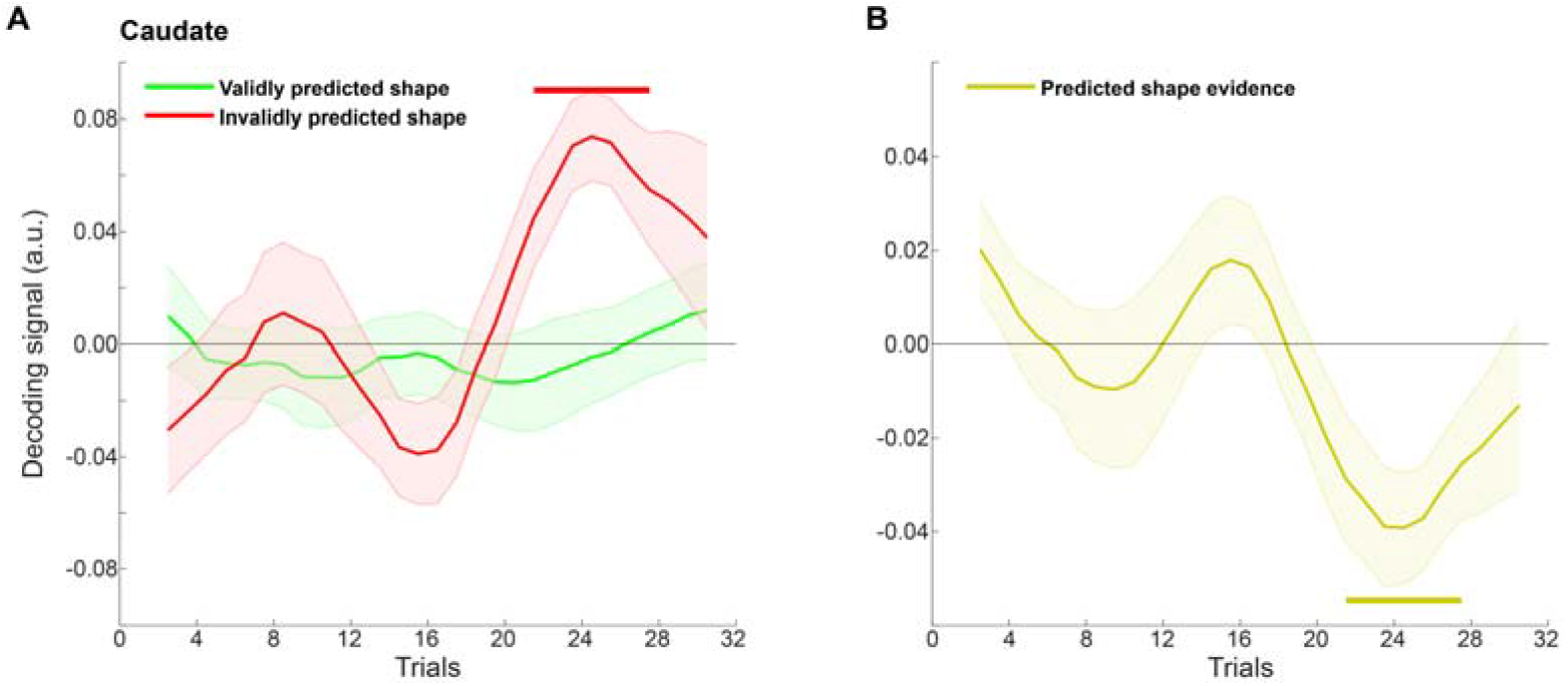
Experiment 1 shape decoding over trials in the caudate nucleus. **A)** Decoding evidence for validly (green) and invalidly (red) predicted shapes in the caudate. **B)** Decoding evidence for predicted (valid – invalid) shapes in the caudate. Horizontal lines indicate significant clusters. Shaded regions indicate SEM.

**Figure S3.**
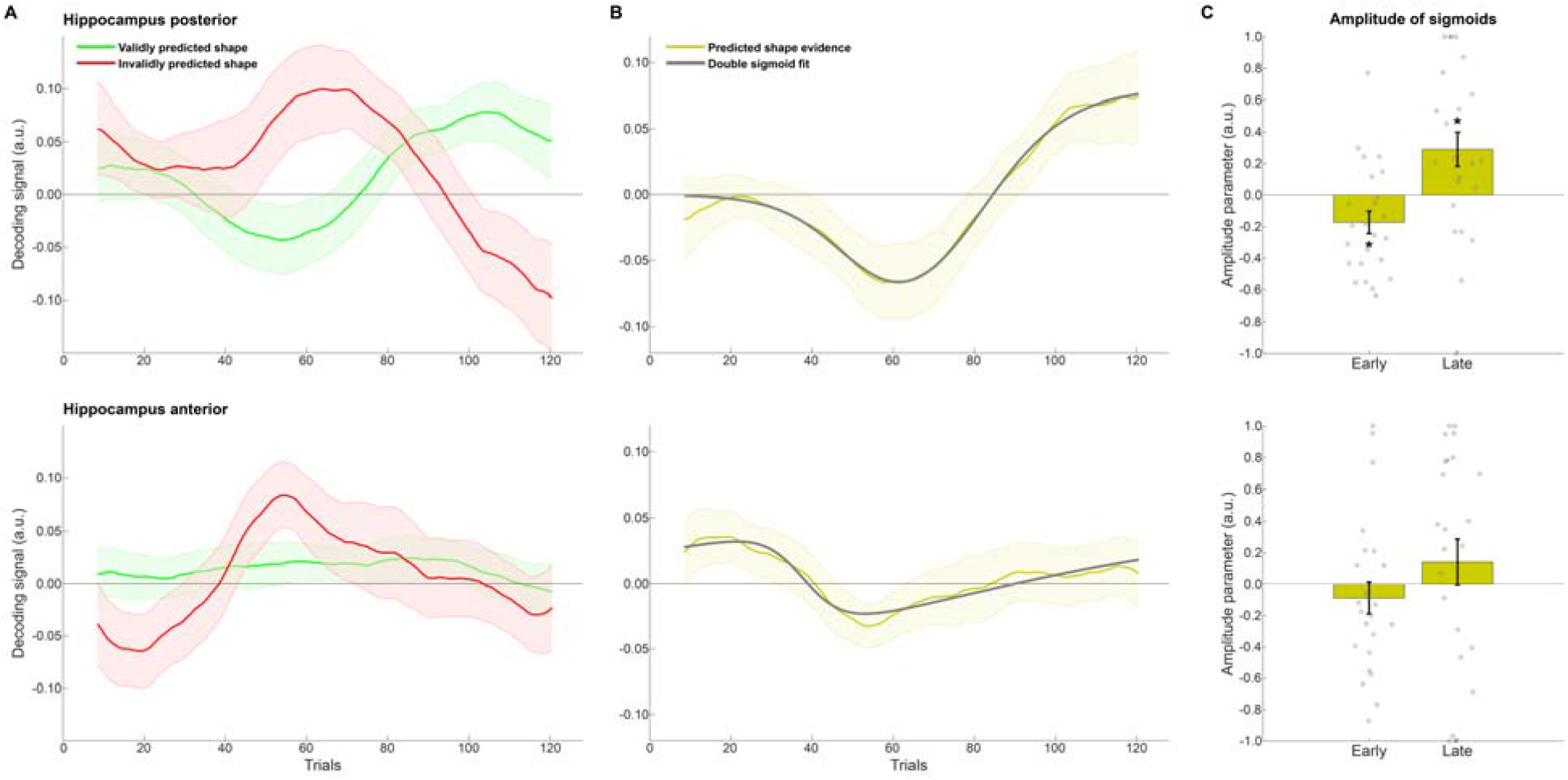
Experiment 2 shape decoding over trials in posterior and anterior hippocampus. **A)** Decoding evidence for validly (green) and invalidly (red) predicted shapes in posterior hippocampus. **B)** Decoding evidence for predicted (valid – invalid) shapes in posterior hippocampus (yellow) with double sigmoid fit (gray). **C)** Amplitude parameters of early (midpoint between trials 1 and 64) and late (midpoint between trials 65 and 128) sigmoid curves in posterior hippocampus. **D)** Decoding evidence for validly (green) and invalidly (red) predicted shapes in anterior hippocampus. **E)** Decoding evidence for predicted (valid – invalid) shapes in anterior hippocampus (yellow) with double sigmoid fit (gray). **F)** Amplitude parameters of early (midpoint between trials 1 and 64) and late (midpoint between trials 65 and 128) sigmoid curves in anterior hippocampus. Shaded regions and error bars indicate SEM. Dots indicate individual participants. *p < 0.05.

**Figure S4.**
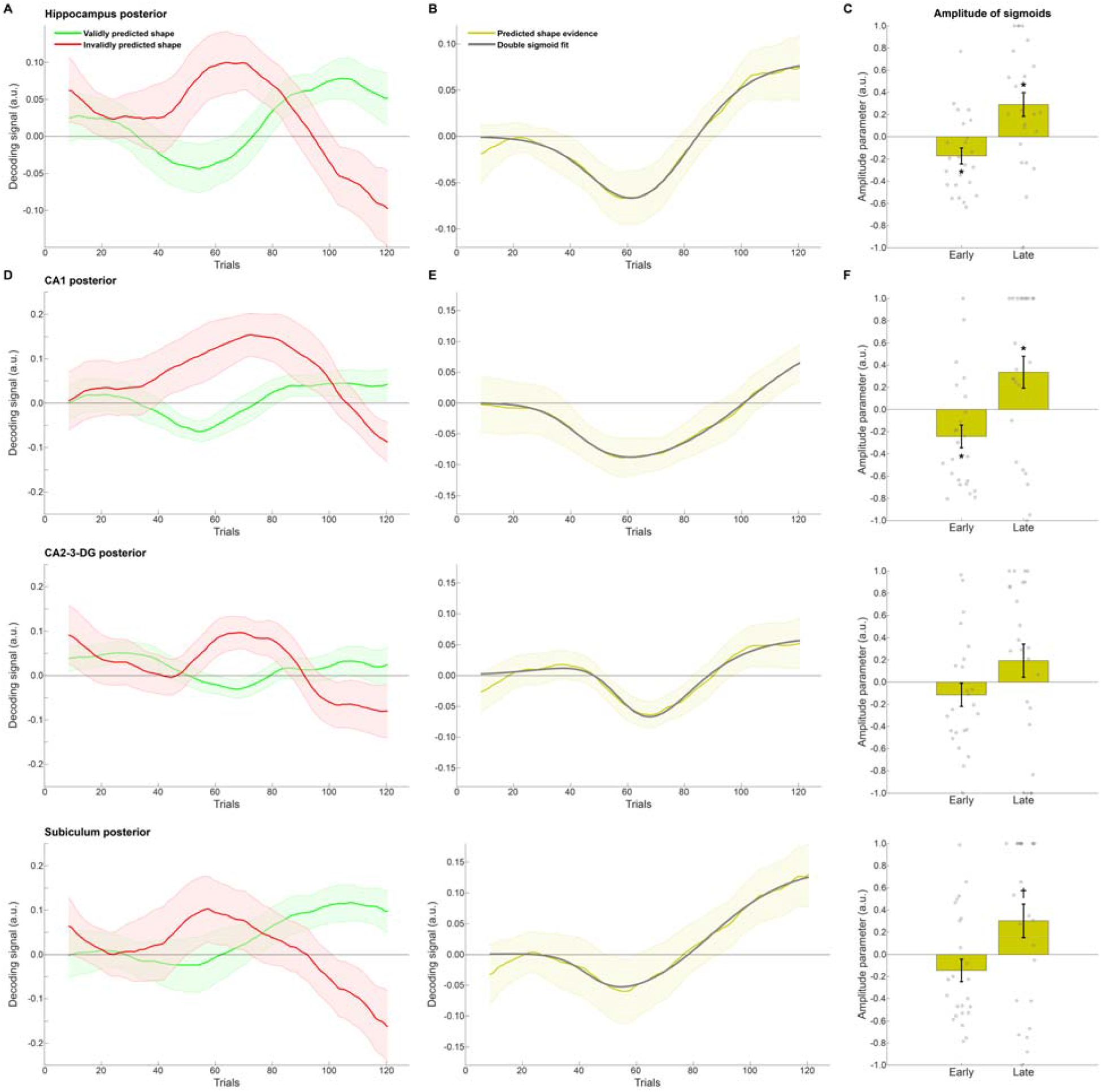
Experiment 2 shape decoding over trials in posterior hippocampus. **A)** Decoding evidence for validly (green) and invalidly (red) predicted shapes in posterior hippocampus. **B)** Decoding evidence for predicted (valid – invalid) shapes in posterior hippocampus (yellow) with double sigmoid fit (gray). **C)** Amplitude parameters of early (midpoint between trials 1 and 64) and late (midpoint between trials 65 and 128) sigmoid curves in hippocampus. **D)** Decoding evidence for validly (green) and invalidly (red) predicted shapes in posterior hippocampal subfields. **E)** Decoding evidence for predicted (valid – invalid) shapes in posterior hippocampal subfields (yellow) with double sigmoid fit (gray). **F)** Amplitude parameters of early (midpoint between trials 1 and 64) and late (midpoint between trials 65 and 128) sigmoid curves in posterior hippocampal subfields. Shaded regions and error bars indicate SEM. Dots indicate individual participants. *p < 0.05; ^†^p = 0.055.

**Figure S5.**
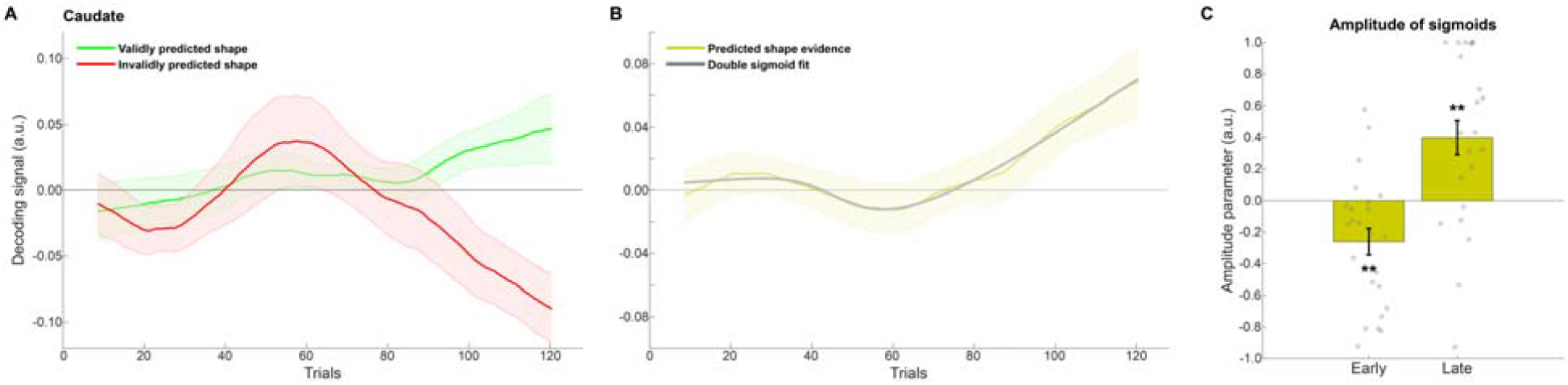
Experiment 2 shape decoding over trials in the caudate nucleus. **A)** Decoding evidence for validly (green) and invalidly (red) predicted shapes in the caudate. **B)** Decoding evidence for predicted (valid – invalid) shapes in the caudate (yellow) with double sigmoid fit (gray). **C)** Amplitude parameters of early (midpoint between trials 1 and 64) and late (midpoint between trials 65 and 128) sigmoid curves in the caudate. Shaded regions and error bars indicate SEM. Dots indicate individual participants. **p < 0.01.

